# Glucocorticoid signaling in pancreatic islets modulates gene regulatory programs and genetic risk of type 2 diabetes

**DOI:** 10.1101/2020.05.15.038679

**Authors:** Anthony Aylward, Mei-Lin Okino, Paola Benaglio, Joshua Chiou, Elisha Beebe, Jose Andres Padilla, Sharlene Diep, Kyle J Gaulton

**Author notes:** Corresponding author: Kyle J Gaulton, 9500 Gilman Drive, #0746, Department of Pediatrics, University of California San Diego. Authors contributed equally to this work (Anthony Aylward, Mei-Lin Okino).

## Abstract

Glucocorticoids are key regulators of glucose homeostasis and pancreatic islet function, but the gene regulatory programs driving responses to glucocorticoid signaling in islets and the contribution of these programs to diabetes risk are unknown. In this study we used ATAC-seq and RNA-seq to map chromatin accessibility and gene expression from eight primary human islet samples cultured *in vitro* with the glucocorticoid dexamethasone. We identified 2,838 accessible chromatin sites and 1,114 genes with significant changes in activity in response to glucocorticoids. Chromatin sites up-regulated in glucocorticoid signaling were prominently enriched for glucocorticoid receptor binding sites and up-regulated genes were enriched for ion transport and lipid metabolism, whereas down-regulated chromatin sites and genes were enriched for inflammatory, stress response and proliferative processes. Genetic variants associated with glucose levels and T2D risk were enriched in glucocorticoid-responsive chromatin sites, including fine-mapped risk variants at 54 known signals. Among fine-mapped variants in glucocorticoid-responsive chromatin, a likely casual variant at the 2p21 locus had glucocorticoid-dependent allelic effects on beta cell enhancer activity and affected *SIX2* and *SIX3* expression. Our results provide a comprehensive map of islet regulatory programs in response to glucocorticoids through which we uncover a role for islet glucocorticoid signaling in mediating risk of type 2 diabetes.

## Introduction

Glucocorticoids are steroid hormones produced by the adrenal cortex which broadly regulate inflammatory, metabolic and stress responses and are widely used in the treatment of immune disorders^1–3^. The metabolic consequences of glucocorticoid action are directly relevant to diabetes pathogenesis, as chronic glucocorticoid exposure causes hyperglycemia and steroid-induced diabetes and endogenous excess of glucocorticoids causes Cushing’s syndrome in which diabetes is a common co-morbidity^4,5^. Glucocorticoids contribute to the development of diabetes both through insulin resistance and obesity via effects on adipose, liver and muscle, as well as through pancreatic islet dysfunction^4^. In islets, glucocorticoid signaling has been shown to modulate numerous processes such as insulin secretion, ion channel activity, cAMP signaling, proliferation and development^6–11^.

The effects of glucocorticoids on cellular function are largely mediated through regulation of transcriptional activity. Glucocorticoids diffuse through the cell membrane into cytoplasm and bind the glucocorticoid receptor (GR), which is then translocated into the nucleus where it binds DNA and modulates the transcriptional program^12–15^. Gene activity can be affected by GR via direct genomic binding and regulation as well as indirectly through physical interaction with other transcriptional regulators^13–17^. Previous studies have profiled glucocorticoid signaling by mapping genomic locations of GR binding and other epigenomic features such as histone modifications and chromatin accessibility in response to endogenous glucocorticoids such as cortisol or analogs such as dexamethasone^13,14,18,19^. Studies have also shown that the genomic function of GR is largely mediated via binding to regions of accessible chromatin^20,21^.

Genetic studies have identified hundreds of genomic loci that contribute to diabetes risk and which primarily map to non-coding sequence and affect gene regulation^22–25^. Risk variants for type 2 diabetes (T2D) are enriched for pancreatic islet regulatory sites^22–24,26,27^, while type 1 diabetes (T1D) risk variants are enriched for immune cell as well as islet regulatory sites. The specific mechanisms of most risk variants in islets are unknown, however, which is critical for understanding the genes and pathways involved in disease pathogenesis and for the development of novel therapeutic strategies. Previous studies of islet chromatin have focused predominantly on normal, non-disease states^27,31–36^, although recent evidence has shown that diabetes risk variants can interact with environmental stimuli to affect islet chromatin and gene regulatory programs^30^.

The effects of glucocorticoid and other steroid hormone signaling on islet regulatory programs and how these signals interact with diabetes risk variants, however, are largely unknown. In this study we profiled islet accessible chromatin and gene expression in primary human pancreatic islets exposed *in vitro* to the glucocorticoid dexamethasone. Glucocorticoid signaling had widespread effects on islet accessible chromatin and gene expression levels. Up-regulated chromatin sites were strongly enriched for glucocorticoid receptor binding and up-regulated genes were enriched for processes related to ion channel activity and steroid and lipid metabolism. Conversely, down-regulated sites and genes were involved in inflammation, stress response and proliferation. Genetic variants affecting T2D risk and glucose levels were significantly enriched in glucocorticoid-responsive chromatin sites, including a likely causal variant at the *SIX2/3* locus which had glucocorticoid-dependent effects on beta cell enhancer activity and affected *SIX2* and *SIX3* expression. Together our results provide a comprehensive map of islet gene regulatory programs in response to glucocorticoids which will facilitate a greater mechanistic understanding of glucocorticoid signaling and its role in islet function and diabetes risk.

## Results

### Map of gene regulation in pancreatic islets in response to glucocorticoid signaling

In order to determine the effects of glucocorticoid signaling on pancreatic islet regulation, we cultured primary islet cells *in vitro* with dexamethasone (100 ng/mL for 24hr) as well as in untreated conditions and measured accessible chromatin and gene expression levels in both treated and untreated cells. An overview of the study design is provided in **Figure 1A**.

**Figure 1.**
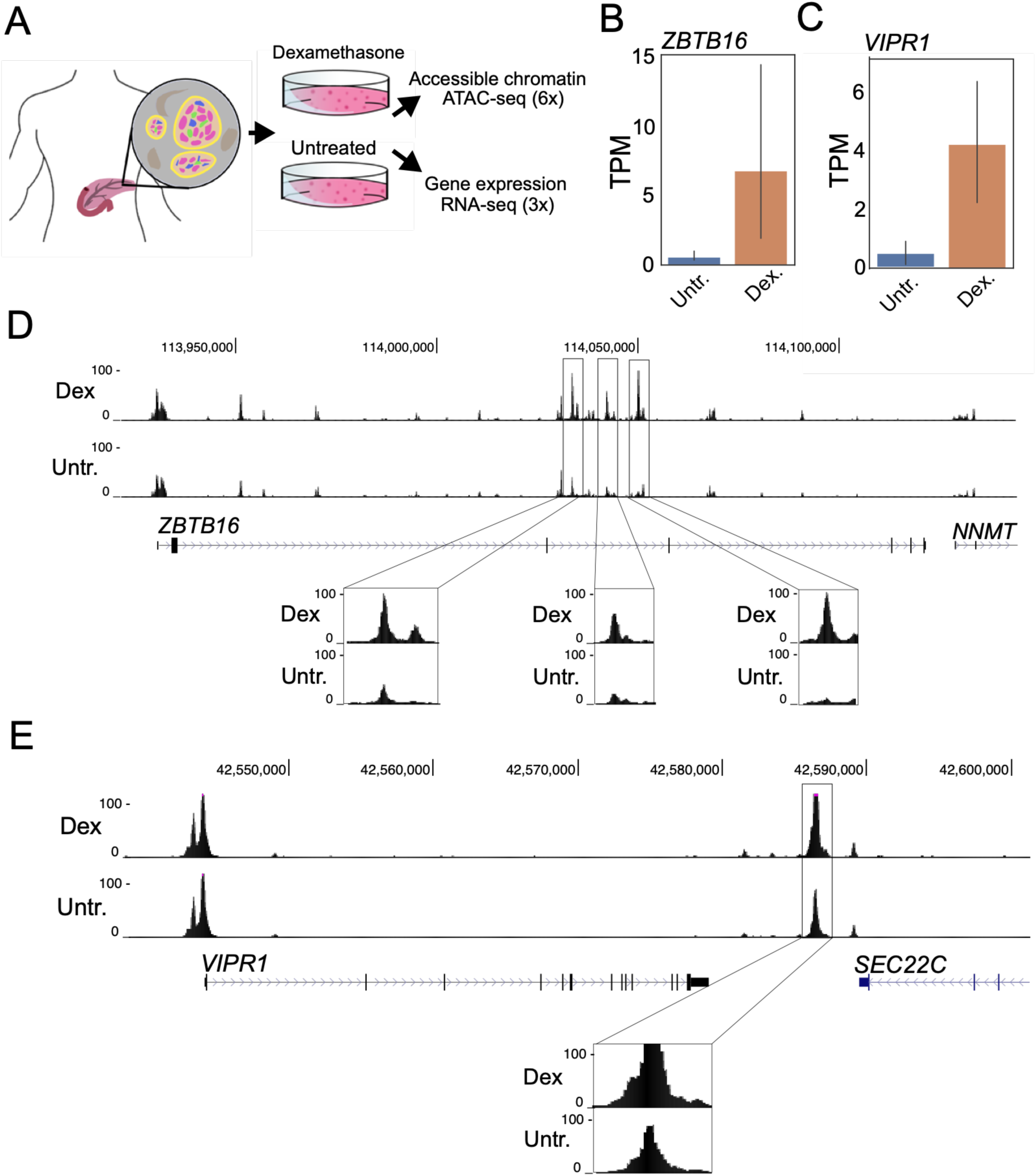
A map of gene regulation in pancreatic islets in response to glucocorticoid signaling. (A) Overview of study design. Primary pancreatic islet samples were split and separately cultured in normal conditions and including the glucocorticoid dexamethasone, and then profiled for gene expression and accessible chromatin using RNA-seq and ATAC-seq assays. (B,C) Genes with known induction in glucocorticoid signaling *ZBTB16* and *VIPR1* had increased expression levels in glucocorticoid-treated islets compared to untreated islets. TPM = transcripts per million. (C) At the *ZBTB16* locus several accessible chromatin sites intronic to *ZBTB16* had increased accessibility in glucocorticoid treated (Dex.) compared to untreated (Untr.) islets. (D) At the *VIPR1* locus an accessible chromatin site downstream of *VIPR1* had increased accessibility in glucocorticoid treated (Dex.) compared to untreated (Untr.) islets. Values in C and D represent RPKM normalized ATAC-seq read counts.

We assayed gene expression in dexamethasone-treated and untreated islets from 3 samples using RNA-seq (**Supplemental Table 1; see Methods**). Across replicate samples we observed changes in expression levels of genes both known to be induced by dexamethasone such as *ZBTB16*^37–39^ and *VIPR1*^40^ as well as those suppressed by dexamethasone such as *IL11*^41^ (**Figure 1B, Figure 1C, Supplemental Figure 1A**). We next assayed accessible chromatin in dexamethasone-treated and untreated islets from 6 samples using ATAC-seq (**Supplemental Table 1; see Methods**). Across replicate samples we observed reproducible changes in islet accessible chromatin signal concordant with changes in gene expression. For example, accessible chromatin signal was notably induced at several sites proximal to the *ZBTB16* and *VIPR1* genes in dexamethasone-treated compared to untreated islets (**Figure 1B**,**C, Supplemental Figure 2, Supplemental Figure 3**). Similarly, accessible chromatin signal was reduced at sites proximal to *IL11* in glucocorticoid-treated compared to untreated islets (**Supplemental Figure 1B**).

**Figure 2.**
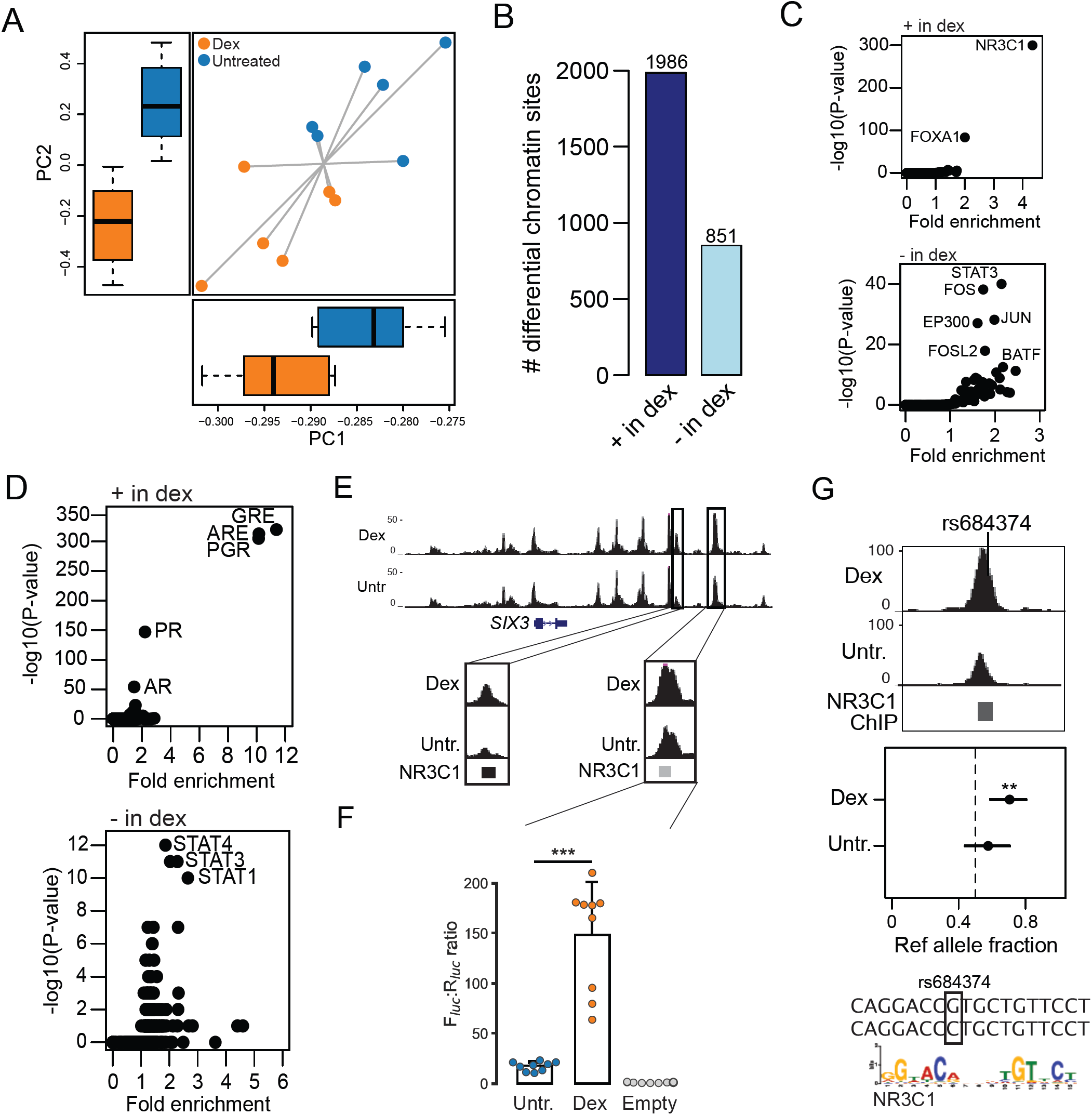
Glucocorticoid signaling affects chromatin accessibility in pancreatic islets. (A) Principal components plot showing ATAC-seq signal for 6x glucocorticoid-treated (orange) and untreated (blue) islets. Lines connect paired assays from the same sample, and box plots on each axis represent the average values for each condition. (B) Number of sites with differential chromatin accessibility in glucocorticoid treated compared to untreated islets, including sites with increased activity (+ in dex) and decreased activity (- in dex). (C) Enrichment of ChIP-seq sites from ENCODE for 160 TFs in differential chromatin sites with increased activity (top, + in dex) and decreased activity (bottom, - in dex) in glucocorticoid treated islets. (D) Sequence motifs enriched in differential chromatin sites with increased activity (top, + in dex) and decreased activity (bottom, - in dex) in glucocorticoid-treated islets. (E) Multiple chromatin sites at the *SIX2/3* locus had increased activity in glucocorticoid-treated islets and overlapped ChIP-seq sites for the glucocorticoid receptor (GR/NR3C1) (top). (F) One of the differential sites at *SIX2/3* had glucocorticoid-dependent effects on enhancer activity in gene reporter assays in MIN6 cells (bottom). Values represent mean and standard deviation. ***P=1.6×10^−6^. (G) Variant rs684374 mapped in a chromatin site with increased activity in glucocorticoid treated islets, had significant allelic effects on chromatin accessibility specifically in glucocorticoid-treated islets, and also disrupted a sequence motif for the glucocorticoid receptor. **P=3.8×10-4.

**Figure 3.**
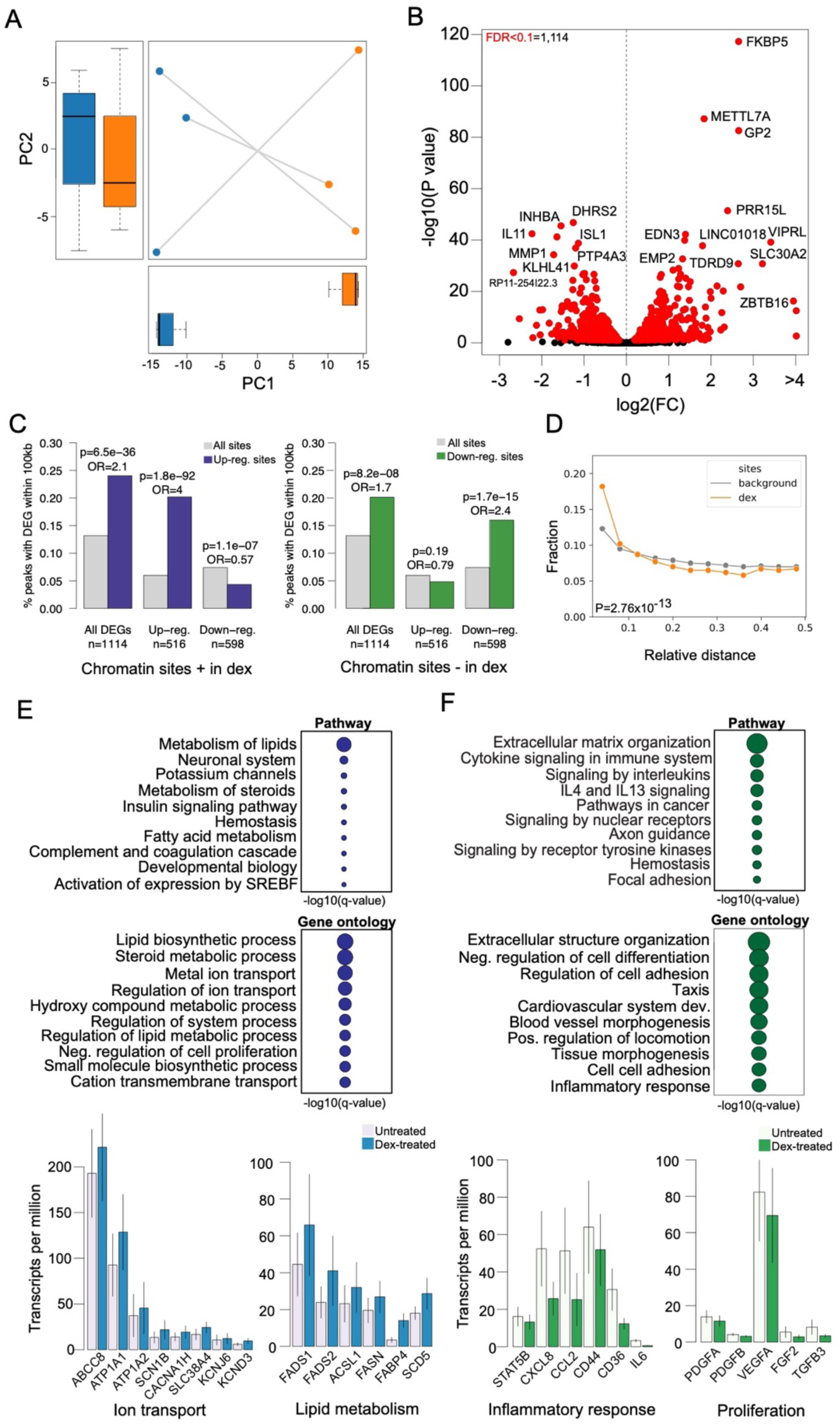
Glucocorticoid signaling affects gene expression levels in pancreatic islets. (A) Principal components plot showing RNA-seq signal for 3x glucocorticoid-treated (orange) and untreated (blue) islets. Lines connect paired assays from the same sample, and box plots on each axis represent the average values for each condition. (B) Volcano plot showing genes with differential expression in glucocorticoid-treated islets compared to untreated islets. Genes with significantly differential expression (FDR<.10) are highlighted in red, and genes with pronounced changed in expression are listed. (C) Percentage of chromatin sites with increased activity (left) and decreased activity (right) in glucocorticoid-treated islets within 100kb of differentially expressed genes compared to chromatin sites without differential activity. (D) Relative distance of accessible chromatin sites with differential activity (dex) to genes with differential expression compared to all chromatin sites (background). (E) Biological pathway and Gene Ontology terms enriched among genes with up-regulated expression in glucocorticoid-treated islets (top), and the expression level of selected genes annotated with ion transport and lipid metabolism terms in glucocorticoid-treated and untreated islets (bottom). (F) Biological pathway and Gene Ontology terms enriched among genes with up-regulated expression in glucocorticoid-treated islets (top), and the expression level of selected genes annotated with inflammatory response and proliferation terms in glucocorticoid-treated and untreated islets (bottom). Circles represent - log10 of the enrichment q-value, and bar plots represent mean and standard error.

### Islet accessible chromatin sites with differential activity in response to glucocorticoid signaling

To understand the effects of glucocorticoid signaling on accessible chromatin in islets at a genome-wide level, we first performed principal components analysis (PCA) using the read counts in chromatin sites for each treated and untreated islet ATAC-seq sample (**see Methods**). We observed reproducible differences in accessible chromatin profiles in dexamethasone-treated islets compared to untreated islets across replicate samples (**Figure 2A**).

We then identified specific islet accessible chromatin sites with significant differential activity in glucocorticoid treatment compared to untreated control cells (**see Methods**). We observed 2,838 sites genome-wide with significant evidence (FDR<.10) for differential activity in glucocorticoid signaling (**Figure 2B, Supplemental Table 2**). Among these 2,838 glucocorticoid-responsive sites, 1,986 had up-regulated activity and 851 had down-regulated activity in glucocorticoid treated compared to untreated cells (**Figure 2B, Supplemental Table 2**). The majority of sites (95%) with differential activity were already accessible in untreated islets, suggesting that sites induced by glucocorticoid signaling typically not activated *de novo*. Furthermore, a majority of differentially accessible sites (2,500, 88%) were not proximal to promoter regions, suggesting they act via distal regulation of gene activity.

We next characterized transcriptional regulators underlying changes in glucocorticoid-responsive islet chromatin. First, we identified TF motifs enriched in genomic sequence underneath sites up-regulated and down-regulated in glucocorticoid-treated islets (**see Methods**). The most enriched sequence motifs in up-regulated sites were for glucocorticoid and other steroid hormone response elements (GRE P<1×10^−300^, ARE P=1×10^−313^, PGR P=1×10^−305^), in addition to lesser enrichment for TFs relevant to islet function (e.g. FOXA1 P=1×10^−11^) (**Figure 2C, Supplemental Table 3)**. Conversely, down-regulated sites were most enriched for sequence motifs for STAT TFs (STAT4 P=1×10^−12^, STAT3 P=1×10^−11^) followed by TFs involved in islet function (NKX6.1 P=1×10^−7^, FOXA2 P=1×10^−6^) (**Figure 2C, Supplemental Table 3**). Next, we determined enrichment of glucocorticoid-responsive chromatin sites for ChIP-seq TF-binding sites previously identified by the ENCODE project. We observed strongest enrichment of up-regulated accessible chromatin sites for glucocorticoid receptor (NR3C1) binding sites (ratio=73.1, P<1×10^−300^), and less pronounced enrichment for binding sites of FOXA1 and other TFs (**Figure 2D, Supplemental Table 3**). Down-regulated sites were most enriched for STAT binding (STAT3 ratio=2.1, P=7.6×10^−41^) as well as enhancer binding TFs such as FOS/JUN and P300 (**Figure 2D, Supplemental Table 3**).

Accessible chromatin sites with significant up-regulation in glucocorticoid signaling compared to untreated islets included several that mapped to the *SIX2/SIX3* locus (**Figure 2E**), which also harbors genetic variants associated with fasting glucose level and risk of T2D. Glucocorticoid-responsive sites at this locus also directly overlapped NR3C1 ChIP-seq sites identified by the ENCODE project (**Figure 2E**). We tested one of the sites up-regulated by glucocorticoids at this locus (fold-change=1.75; P=3.6×10^−5^, **Supplemental Table 2**) for enhancer activity in luciferase gene reporter assays in dexamethasone-treated and untreated MIN6 cells. We observed a significant increase in enhancer activity in dexamethasone-treated cells relative to untreated cells (P=1.65×10^−6^) (**Figure 2F**), confirming that this site is highly induced by glucocorticoid signaling.

We determined the effects of genetic variants on islet accessible chromatin using allelic imbalance mapping. We performed microarray genotyping of islet samples and imputed genotypes into 39M variants (**see Methods**). For variants overlapping islet chromatin sites we obtained read counts in samples heterozygote for that variant, corrected for mapping bias using WASP and modeled the resulting counts for imbalance using a binomial test. We then identified variants with evidence for allelic imbalance (FDR<.10) in accessible chromatin for either glucocorticoid-treated or untreated islets (**Supplemental Table 4**). Among imbalanced variants, several both mapped in glucocorticoid-responsive chromatin and had significant effects in glucocorticoid-treated islets, suggesting that their effects might interact with glucocorticoid signaling. For example, variant rs684374 at 15q14 in a glucocorticoid-responsive site bound by GR had significant imbalance in glucocorticoid-treated islets only (GC P=2.6×10^−4^, untr. P=.22) and was predicted to alter binding of GR (NR3C1) (**Figure 2H**). Similarly, variant rs11610384 at 12p11 in a glucocorticoid-responsive site bound by GR had significant imbalance in glucocorticoid-treated islets only (GC P=1.5×10^−5^, untr. P=1) and disrupted nuclear receptor motifs (**Supplemental Table 4**).

These results demonstrate that glucocorticoid signaling broadly affects accessible chromatin in islets including sites both up-regulated through glucocorticoid receptor activity and down-regulated through the activity of STAT and other TFs.

### Genes and pathways with differential regulation in islets in response to glucocorticoid signaling

We next sought to determine the effects of glucocorticoid treatment on gene expression levels. We first performed PCA using gene transcript counts from untreated and dexamethasone-treated islet samples obtained from RNA-seq assays (**see Methods**). There were again reproducible differences in expression levels across replicate samples (**Figure 3A)**.

We next identified specific genes with differential expression in response to glucocorticoids compared to untreated islet samples using DESeq2 (**see Methods**). There were 1,114 genes with significant evidence for differential expression (FDR<0.10) in glucocorticoid signaling (**Supplemental Table 5**). Among these genes, 46% were up-regulated and 54% were down-regulated in response to glucocorticoids compared to untreated islets (**Figure 3B)**. Genes with the most significant up-regulation included *FBKP5* (log2(FC)=2.65, FDR=4.97×10^−118^), a chaperone of the glucocorticoid receptor, *METTL7A* (log2(FC)=1.84, FDR=6.09×10^−88^), *GP2* (log2(FC)=2.65, FDR=2.48×10^−83^), *PRR15L* (log2(FC)=2.39, FDR=3.73×10^−52^), and *EDN3* (log2(FC)=1.37, FDR=1.19×10^−40^). Conversely, genes with most significant down-regulation included *INHBA* (log2(FC)=-1.54, FDR=2.79×10^−46^), *DHRS2* (log2(FC)=-1.255, FDR=1.51×10^−47^) and *IL11* (log2(FC)=-2.23, FDR=3.46×10^−43^) (**Figure 3B**).

We determined whether changes in gene expression in glucocorticoid signaling were driven through accessible chromatin, by testing for enrichment of glucocorticoid-responsive chromatin sites for proximity to genes with glucocorticoid-responsive changes in expression. Glucocorticoid-responsive chromatin sites were significantly more likely to map within 100kb of a gene with glucocorticoid-responsive expression compared to other chromatin sites in islets (OR=2.0, P=9×10^−41^). We next performed these analyses separately for sites up- and down-regulated in glucocorticoid signaling. There was significant enrichment of sites with increased activity in glucocorticoid signaling within 100kb of up-regulated genes specifically (up OR=4.0, P=1.8×10^−92^; down OR=0.57, P=1.1×10^−7^) (**Figure 3C**). Similarly, sites with decreased activity in glucocorticoid signaling were enriched within 100kb of down-regulated genes (down OR=2.4, P=1.7×10^−15^; up OR=0.79, P=0.19) (**Figure 3C**). Furthermore, we also observed an enrichment of glucocorticoid-responsive chromatin sites for closer proximity to genes with glucocorticoid-responsive expression compared to background sites (Kolmogorov-Smirnov P=2.76×10^−13^) (**Figure 3D**).

In order to understand the molecular pathways affected by glucocorticoid activity in islets, we tested genes up- and down-regulated in glucocorticoid signaling for gene set enrichment using gene ontology (GO) terms (**see Methods**). Up-regulated genes showed strongest enrichment for GO terms related to steroid metabolism (steroid metabolic process P=5.8×10^−20^), and were also enriched for potassium and other ion transport (potassium channels P=7.1×10^−10^; regulation of ion transport P=6.8×10^−18^), lipid metabolism (lipid biosynthetic process P=1.6×10^−20^), and insulin signaling (insulin signaling pathway P=4.5×10^−9^) **(Figure 3E, Supplemental Table 6**). Numerous genes that function in ion transport were up-regulated in glucocorticoid signaling; for example *ATP1A1, ATP2A2, SCN1B, SCNN1A, CACNA1H, CACNG4, SLC38A4, TRPV6* as well as 13 potassium channel genes including *ABCC8, KCNJ2, KCNJ6*, and *KCND3* (**Figure 3E, Supplemental Table 5-6**). Up-regulated genes also included numerous that function in lipid metabolism including *FADS1, FADS2, ACSL1, SCD5, FASN, FABP4, ACACB*, and *ANGPTL4* (**Figure 3E, Supplemental Table 5-6**).

Conversely, genes down-regulated in glucocorticoid signaling were enriched for inflammatory response (inflammatory response P=7.9×10^−21^; cytokine signaling in immune system P=2.9×10^−^ 18), stress response (cellular responses to stress P=3.8×10^−10^), extracellular matrix, cell adhesion and morphogenesis (extracellular matrix organization P=2.3×10^−26^, cell adhesion P=1.2×10^−26^), and cell differentiation and proliferation terms (neg. regulation of cell differentiation P=2.9×10^−27^) (**Figure 3F, Supplemental Table 5-6)**. Down-regulated genes included those involved in the inflammatory response such as *IL6, STAT5B, STAT3, STAT4, SMAD3, CXCL8, STAT3, CCL2, CD44, CD36, RELB, IRF1*, extracellular matrix formation such matrix metalloproteinase genes such as *MMP1, MMP9* and matrix components such as *FBN1*, pancreatic differentiation such as *ISL1, PAX6, NKX6-1, HES1* and *JAG1*, and proliferation and growth factors such as *PDGFA, PDGFB, FGF2, TGFB3* and *VEGFA* (**Figure 3F, Supplemental Table 5-6**).

These results demonstrate that glucocorticoid signaling in islets up-regulates genes involved in steroid metabolism, lipid metabolism and ion channel activity, and down-regulates genes involved in inflammation, stress response, differentiation, proliferation and extracellular matrix formation.

### Enrichment of T2D and glucose associated variants in glucocorticoid-responsive islet chromatin

Genetic variants associated with diabetes risk are enriched in pancreatic islet regulatory elements. As these studies have been performed primarily using non-diabetic donors in normal (untreated) conditions, however, the role of environmental stimuli in modulating diabetes-relevant genetic effects on islet chromatin is largely unknown. We therefore tested diabetes and fasting glycemia associated variants for enrichment in glucocorticoid-responsive islet chromatin sites compared to a background of other islet chromatin sites (**see Methods**). We observed significant enrichment of variants influencing T2D risk and blood sugar (glucose) levels in glucocorticoid-responsive chromatin (T2D -log10(P)=1.35, glucose -log10(P)=1.50) (**Figure 4A**). Conversely, we observed no evidence for enrichment of T1D risk variants (-log10(P)=0.22) (**Figure 4A**).

**Figure 4.**
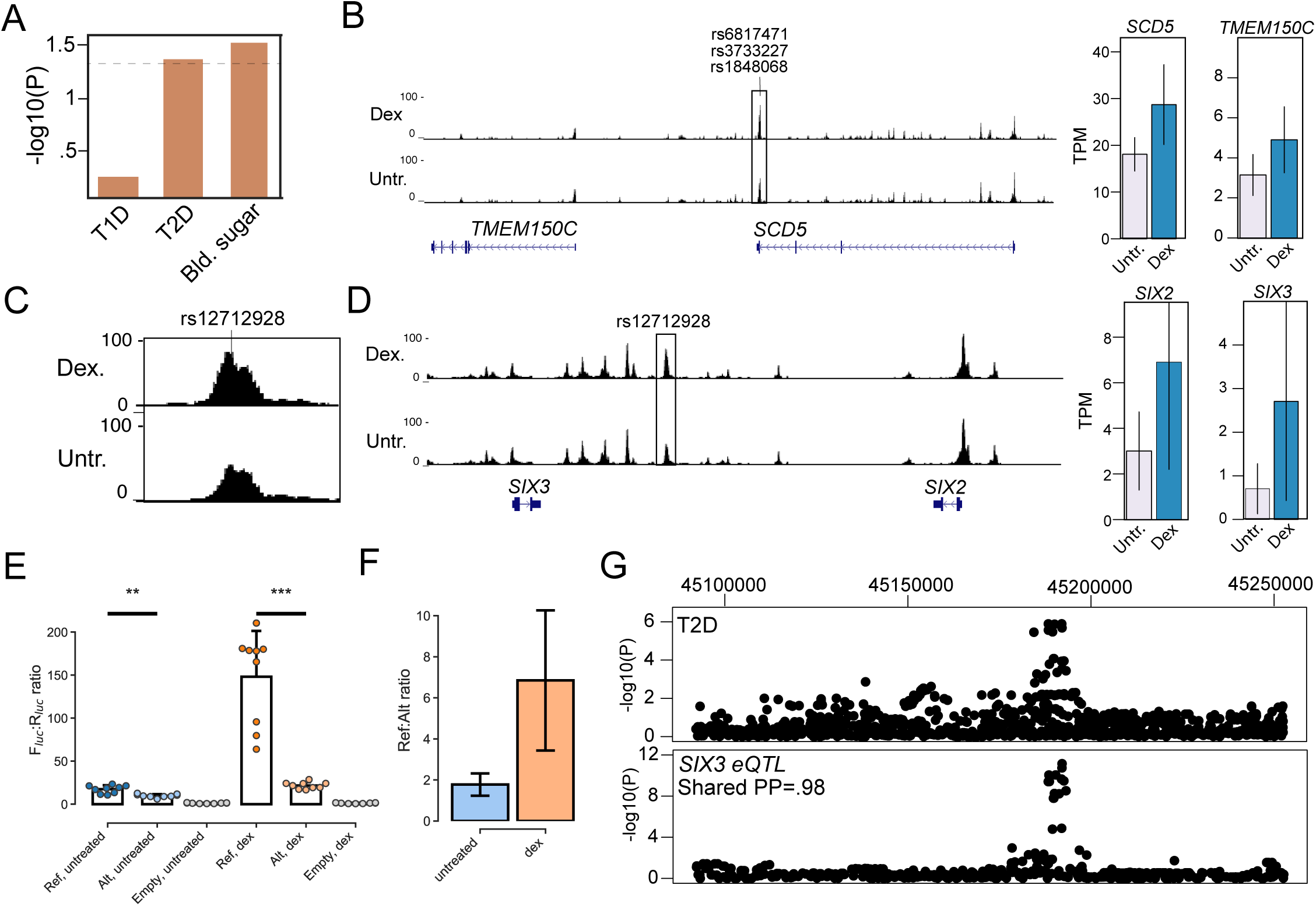
Type 2 diabetes and glucose associated variants affect glucocorticoid-responsive islet regulatory programs. (A) Enrichment of variants associated with type 1 diabetes, type 2 diabetes and blood sugar (glucose) levels for sites with differential chromatin accessibility in glucocorticoid-treated islets. (B) Multiple fine-mapped T2D variants at the *SCD5/TMEM150C* locus mapped in a glucocorticoid-responsive islet accessible chromatin site. Both the *SCD5* and *TMEM150C* genes had increased expression in glucocorticoid-treated islets. TPM = transcripts per million. Genome browser tracks represent RPKM normalized ATAC-seq signal, and TPM bar plots represent mean and standard error. (C, D) Variant rs12712928 with evidence for blood sugar and T2D association mapped in a glucocorticoid-responsive chromatin site at the *SIX2/3* locus. Both the *SIX2* and *SIX3* genes had increased expression in glucocorticoid-treated islets. (E) Variant rs12712928 had significant allelic effects on enhancer activity in gene reporter assays in MIN6 cells. Values represent mean and standard deviation. **P=3.2×10^−4^; ***P=2.5×10^−6^ (F) The allelic effects of rs12712928 were more pronounced in glucocorticoid-treated relative to untreated islets. Values represent fold-change and 95% CI. (G) The T2D association signal at *SIX2/3* was colocalized with an eQTL for *SIX3* expression in islets.

We catalogued fine-mapped variants overlapping glucocorticoid-responsive islet chromatin using 99% credible sets of T2D and glucose level signals from DIAMANTE and Biobank Japan^22,42^ (**see Methods**). We identified 54 signals where a fine-mapped variant overlapped at least one glucocorticoid-responsive site (**Supplemental Table 7)**. We further cataloged 412 variants genome-wide in glucocorticoid-responsive sites with nominal evidence for T2D association (P<.005) in DIAMANTE or Biobank Japan GWAS (**Supplemental Table 7**). We next prioritized potential target genes of T2D-associated variants in glucocorticoid-responsive chromatin by identifying genes proximal to these sites and with expression patterns consistent with the activity of the site (**Supplemental Table 7**). For example, T2D-associated variants at the 11q12 locus mapped in a chromatin site induced by glucocorticoid signaling proximal to *SCD5* and *TMEM150C* which both had up-regulated expression (**Figure 4B, Supplemental Table 5, Supplemental Table 7**). Similarly, T2D-associated variants at the 4q31 locus mapped in a chromatin site down-regulated in glucocorticoid signaling proximal to *FBXW7* which had down-regulated expression (**Supplemental Figure 4A, Supplemental Table 5, Supplemental Table 7**). Outside of known T2D loci we observed numerous additional examples such as rs1107376 (T2D P=2.2×10^−4^) in a site induced in glucocorticoids which was proximal to *NPY* which had glucocorticoid-stimulated expression (**Supplemental Figure 4B, Supplemental Table 5, Supplemental Table 7)**.

At the 2p21 locus, glucose level-associated variant rs12712928 mapped in a chromatin site with increased activity in glucocorticoid signaling and was proximal to *SIX2* and *SIX3* which both had glucocorticoid-induced expression (**Figure 4C**,**D, Supplemental Table 5, Supplemental Table 7**). This variant had the highest posterior probability in glucose fine-mapping data (PPA=.89), suggesting it is causal for the association signal at this locus. Furthermore, this variant also had evidence for T2D association (Biobank Japan T2D P=2.1×10^−6^), strongly suggesting that it influences T2D risk as well. We therefore tested whether rs12712928 affected enhancer activity using sequence around variant alleles in untreated and dexamethasone treated MIN6 cells (**see Methods**). The glucose level increasing and T2D risk allele C had significantly reduced enhancer activity in both glucocorticoid-treated (P=2.5×10^−6^) and untreated cells (P=3.2×10^−4^) (**Figure 4E**). However, the allelic differences at this variant were more pronounced in glucocorticoid-treated cells (ref/alt ratio GC=6.85, 95% CI=3.4,10.2; untreated=1.78, 95% CI=1.23,2.32) (**Figure 4F**). We next identified gene(s) directly affected by rs12712928 activity using expression QTL data in islets^35^. We observed evidence that rs10168523 was an islet QTL for *SIX3* and *SIX2* expression (*SIX3* eQTL P=1.8×10^−11^, *SIX2* eQTL P=1.6×10^−6^; **Figure 4G**), where the T2D risk allele was correlated with reduced expression of both genes. Glucose level and T2D association at this locus was also strongly co-localized with both the *SIX3* and *SIX2* eQTLs (T2D shared *SIX3* PP=98%, *SIX2* PP=91%; Blood sugar shared *SIX3* PP=99%, *SIX2* PP=99%) (**Figure 4G**).

These results demonstrate that T2D and glucose level variants are enriched in glucocorticoid-responsive chromatin sites in islets, including variants that interact with glucocorticoid signaling directly to affect islet regulation.

## Discussion

Our study demonstrates the relevance of islet chromatin dynamics in response to corticosteroid signaling to T2D pathogenesis, including T2D risk variants that interact with corticosteroid activity directly to affect islet chromatin. In a similar manner, variants mediating epigenomic responses of pancreatic islets to proinflammatory cytokines were recently shown to contribute to genetic risk of T1D^30^. Numerous environmental signals and external conditions modulate pancreatic islet function and contribute to the pathophysiology and genetic basis of diabetes, yet the epigenomic and transcriptional responses of islets to disease-relevant stimuli have not been extensively measured. Future studies of islet chromatin and gene regulation exposed to additional stimuli will therefore likely continue providing additional insight into diabetes risk.

Glucocorticoid signaling led to widespread changes in accessible chromatin, which up-regulated the expression of proximal genes enriched for processes related to ion channels and transport, in particular potassium channels. Potassium ion concentrations modulate calcium influx and insulin secretion in beta cells^43^, and in disruption of ion channel function leads to impaired glucose-induced insulin secretion and diabetes^44^. Glucocorticoids have been shown to suppress calcium influx while preserving insulin secretion via cAMP^7^, and in line with this finding we observed increased activity of potassium channel and cAMP signaling genes. Up-regulated genes were also enriched in lipid metabolism, which has been shown to regulate insulin secretion and contribute to diabetes^45,46^. Several up-regulated genes *PER1* and *CRY2* are also components of the circadian clock, and previous studies have shown that endogenous glucocorticoid release is under control of circadian rhythms and therefore may contribute to downstream regulation of the clock^47^. Conversely, glucocorticoid signaling down-regulated inflammatory and stress response programs, in line with previous reports and the known function of glucocorticoids^2,17,48^. Our findings further suggest that down-regulation of gene activity in glucocorticoid signaling is mediated through the activity of STAT and other TFs at proximal accessible chromatin sites, either through reduced TF expression or inhibition by GR.

Genetic variants near the homeobox TFs *SIX2* and *SIX3* influence glucose levels^49,50^, and our results provide evidence that both of these TFs operate downstream of glucocorticoid signaling and that the variants interact with this signaling program directly to influence glucose levels and risk of T2D. A previous study identified association between this locus and glucose levels in Chinese samples and demonstrated allelic effects of the same variant on islet enhancer activity and binding of the TF GABP^50^, further supporting the likely causality of this variant. *SIX2* and *SIX3* have been widely studied for their role in forebrain, kidney and other tissue development^51–56^. In islets, *SIX2* and *SIX3* both have been shown to increase expression in adult compared to juvenile islets, and induction of *SIX3* expression in EndoB-CH1 cells and juvenile islets enhanced islet function, insulin content and secretion and may contribute to the suppression of proliferative programs^57^. These findings are in line with those of our study which reveal that corticosteroid signaling increases the activity of genes involved in islet function and insulin secretion while suppressing inflammatory and proliferative gene activity.

Our *in vitro* experimental model mimics the environment of pancreatic islets under hormone signaling, albeit for a single treatment and condition. Given the similarity in binding sites of many nuclear hormone receptors, the effects of GR binding on gene regulation may overlap with the activity of other nuclear receptors which act in beta cells^58^. Studies of other tissues have profiled glucocorticoid signaling across a range of experimental conditions and identified dose- and temporally-dependent effects on gene regulatory programs^14,15^, and in islets dose- and temporally-dependent effects of glucocorticoids may impact insulin secretion and other islet functions. Future studies profiling the genomic activity of nuclear receptors in islets across a breadth of experimental conditions will therefore help further shed light into the role of hormone signaling dynamics in islet gene regulation and diabetes pathogenesis.

## Methods

### Human islet samples

Human islet samples were obtained through the Integrated Islet Distribution Program (IIDP) and University of Alberta. Islet samples were further enriched using a dithizone stain. Islets were cultured at approximately 10mL media/1k islets in 10cm dishes at 37C, 5% CO2 in CMRL 1066 media supplemented with 10% FBS, 1X pen-strep, 8mM glucose, 2mM L-glutamine, 1mM sodium pyruvate, 10mM HEPES, and 250ng/mL Amphotericin B. Treated islets had an additional 100 ng/mL dexamethasone (Sigma) added in the culture media. Islet studies were approved by the Institutional Review Board of the University of California San Diego.

### ATAC-seq assays

Islet samples were collected and centrifuged at 500xg for 3 minutes, then washed twice in HBSS, and resuspended in nuclei permeabilization buffer consisting of 5% BSA, 0.2% IGEPAL-CA630, 1mM DTT, and 1X complete EDTA-free protease inhibitor (Sigma) in 1X PBS. Islets were homogenized using a chilled glass dounce homogenizer and incubated on a tube rotator for 10 mins before being filtered through a 30uM filter (sysmex) and centrifuged at 500xg in a 4C microcentrifuge to pellet nuclei. Nuclei were resuspended in Tagmentation Buffer (Illumina) and counted using a Countess II Automated Cell Counter (Thermo). Approximately 50,000 nuclei were transferred to a 0.2mL PCR tube and volume was adjusted to 22.5uL with Tagmentation Buffer. 2.5uL TDE1 (Illumina) was added to each tagmentation reaction and mixed with gentle pipetting. Transposition reactions were incubated at 37C for 30 minutes. Tagmentation reactions were cleaned up using 2X reaction volume of Ampure XP beads (Beckman Coulter) and eluted in 20uL Buffer EB (Qiagen). 10uL tagmented DNA prepared as described above was used in a 25uL PCR reaction using NEBNext High-Fidelity Master Mix (New England Biolabs) and Nextera XT Dual-Indexed primers (Nextera). Final libraries were double size selected using Ampure XP beads and eluted in a final volume of 20uL Buffer EB. Libraries were analyzed using the Qubit HS DNA assay (Thermo) and Agilent 2200 Bioanalyzer (Agilent Biotechnologies). Libraries were sequenced on an Illumina HiSeq 4000 using paired end reads of 100bp.

### RNA-seq assays

RNA was isolated from treated and untreated islets using RNeasy Mini kit (Qiagen) and submitted to the UCSD Institute for Genomic Medicine to prepare and sequence ribodepleted RNA libraries. Libraries were sequenced on an Illumina HiSeq4000 using paired end reads of 100bp.

### ATAC-seq data processing

We trimmed reads using Trim Galore with options ‘–paired’ and ‘–quality 10’, then aligned them to the hg19 reference genome using BWA^59^ mem with the ‘-M’ flag. We then used samtools^60^ to fix mate pairs, sort and index read alignments, used Picard (http://broadinstitute.github.io/picard/) to mark duplicate reads, and used samtools^60^ to filer reads with flags ‘-q 30’, ‘-f 3’, ‘-F 3332’. We then calculated the percentage of mitochondrial reads and percentage of reads mapping to blacklisted regions and removed all mitochondrial reads. Peaks were called using MACS2^61^ with parameters ‘—extsize 200 –keep-dup all –shift -100 –nomodel’. We calculated a TSS enrichment score for each ATAC-seq experiment using the Python package ‘tssenrich’. To obtain read depth signal tracks, we used bamCoverage^62^ to obtain bigWig files for each alignment with signal normalization using RPKM.

### Identifying differential chromatin sites

We generated a set of ATAC-seq peaks by merging peaks called from treated and untreated cells across all samples. The set of alignments for each assay were supplied as inputs to the R function featureCounts from the Rsubread^63^ package to generate a read count matrix. We applied the R function DESeqDataSetFromMatrix from the DESeq2^64^ package to the read count matrix with default parameters then applied the DESeq function including donor as a variable to model paired samples. We considered sites differentially accessible with FDR<0.1, as computed by the Benjamini-Hochberg method.

### Principal components analysis

A consensus set of ATAC-seq peaks was defined by merging overlapping (1bp or more) peaks identified in at least two experiments across all ATAC-seq experiments. We constructed a read count matrix using edgeR^65^ and calculated normalization factors using the ‘calcNormFactors’ function. We applied the voom transformation^66^ and used the ‘removeBatchEffect’ function from limma^67^ to regress out batch effects and sample quality effects (using TSS enrichment as a proxy for sample quality). We then restricted the read count matrix to the 10,000 most variable peaks and performed PCA analysis using the core R function ‘prcomp’ with rank 2.

### TF enrichment analysis

Differentially accessible chromatin sites were analyzed for motif enrichment compared to a background of all chromatin sites tested for differential activity using HOMER^68^ and a masked hg19 reference genome with the command ‘findMotifsGenome.pl <bed file> <masked hg19> <output dir> -bg <background bed file> -size 200 -p 8 -bits -preparse -preparsedDir tmp’. For TF ChIP-seq enrichment, we obtained ChIP-seq binding sites for 160 TFs generated by the ENCODE project^69^ and tested for enrichment of binding in differential accessible chromatin sites compared to a background of all remaining chromatin sites genome-wide without differential activity. For each TF we calculated a 2×2 contingency table of overlap with differential sites and non-differential sites, determined significance using a Fisher test and calculated a fold-enrichment of overlap in differential compared to non-differential sites.

### RNA-seq data processing and analysis

Paired-end RNA-Seq reads were aligned to the genome using STAR^70^ (2.5.3a) with a splice junction database built from the Gencode v19 gene annotation^71^. Gene expression values were quantified using the RSEM package (1.3.1) and filtered for >1 TPM on average per sample. Raw expression counts from the remaining 13,826 genes were normalized using variance stabilizing transformation (vst) from DESeq2^64^ and corrected for sample batch effects using limma removeBatchEffect. Principal component analysis was performed in R using the prcomp function. To identify differentially expressed genes between treated and untreated samples we used RSEM^72^ raw expression counts from the 13,826 genes and applied DESeq2^64^ with default settings, including donor as a cofactor to model paired samples. To identify enriched GO terms in up and down-regulated genes, we applied GSEA^73^ to 516 up-regulated and 598 down-regulated genes using Gene Ontology terms and pathway terms. We excluded gene sets with large numbers of genes in enrichment tests.

### Proximity of differential chromatin sites to differentially expressed genes

We calculated the percentage of up- and down-regulated accessible chromatin sites mapping within 100kb of (i) all differentially expressed genes, (ii) up-regulated genes and (iii) down-regulated genes compared to non-differentially accessible sites, and determined the significance and odds ratio using a Fisher exact test. We calculated relative distances using bedtools^74^, with either differential chromatin sites or the “background” of all islet accessible chromatin sites as the “a” argument and differentially expressed genes as the “b” argument. We compared the distribution of relative distances from differential sites to the distribution from background sites using a Kolmogorov-Smirnov test.

### Sample genotyping and imputation

Non-islet tissue was collected for four samples during islet picking and used for genomic DNA extraction using the PureLink genomic DNA kit (Invitrogen). Genotyping was performed using Infinium Omni2.5-8 arrays (Illumina) at the UCSD Institute for Genomic Medicine. We called genotypes using GenomeStudio (v.2.0.4) with default settings. We then used PLINK^75^ to filter out variants with 1) minor allele frequency (MAF) less than 0.01 in the Haplotype Reference Consortium (HRC)^76^ panel r1.1 and 2) ambiguous A/T or G/C alleles with MAF greater than 0.4. For variants that passed these filters, we imputed genotypes into the HRC reference panel r1.1 using the Michigan Imputation Server with minimac4. Post imputation, we removed imputed genotypes with low imputation quality (R2<.3).

### Allelic imbalance mapping

We identified heterozygous variant calls in each sample with read depth of at least 10 in both untreated and treated cells, and then used WASP^77^ to correct for reference mapping bias. We retained variants in each sample where both alleles were identified at least 3 times across untreated and treated cells. We then merged read counts at heterozygous SNPs from all samples in untreated and treated cells separately. We called imbalanced variants from the merged counts using a binomial test, and then calculated q-values from the resulting binomial p-values. We considered variants significant at an FDR<.10.

### Genetic association analysis

We tested glucocorticoid-responsive chromatin sites for enrichment of diabetes associations using fine-mapping data for T1D signals from a prior study^29^, for T2D signals from the DIAMANTE consortium and the Japan Biobank studies^22,42^, and for blood sugar signals from the Japan Biobank study^49^. For the Japan Biobank data, we fine-mapped signals ourselves using GWAS summary statistics for blood sugar and type 2 diabetes. For both traits, we calculated approximate Bayes factors (ABF) for each variant as described previously^78^. We then compiled index variants for each significant locus and defined the set of all credible variants as those in within a 5 Mb window and at least low linkage (r^2^>0.1) in the East Asian subset of 1000 Genomes^79^ with each index. For each locus, we calculated posterior probabilities of associations (PPA) by dividing the variant ABF by the sum of ABF for the locus. We then defined the 99% credible sets by sorting variants by descending PPA and retaining variants adding up to a cumulative probability of 99%.

To test for enrichment, we calculated the cumulative posterior probability of variants overlapping differential sites across all signals. We then defined a background set of ATAC-seq peaks by merging peaks from all ATAC-seq experiments. We estimated an empirical distribution for the total posterior probability using 10,000 random draws of peaks from the background equal in number to the DAC sites. We computed a p-value for each treatment by comparing the total posterior probability within DAC sites to the empirical distribution.

We then cataloged all variants in glucocorticoid-responsive chromatin sites in both fine-mapping data and with nominal association (P<.005) genome-wide. For each variant in glucocorticoid-responsive chromatin, we then identified protein-coding genes in GENCODE v33 with differential expression and where the gene body mapped within 100kb of the variant.

### Expression QTL analyses

We obtained islet expression QTL data from a previous meta-analysis of 230 samples^35^. We extracted variant associations at the *SIX2/SIX3* locus and tested for colocalization between T2D and blood sugar association in the Biobank Japan study and *SIX2* and *SIX3* eQTLs using a Bayesian approach^80^. We considered signals colocalized with shared PP greater than 50%.

### Gene reporter assays

To test for allelic differences in enhancer activity at the *SIX2/3* locus, we cloned human DNA sequences (Coriell) containing the reference allele upstream of the minimal promoter in the luciferase reporter vector pGL4.23 (Promega) using the enzymes Sac I and Kpn I. A construct containing the alternate allele was then created using the NEB Q5 SDM kit (New England Biolabs). The primer sequences used were as follows:

Cloning FWD AGCTAGGTACCCCTCATCTGCCTTTCTGGAC

Cloning REV TAACTGAGCTCCAGTGGGTATTGCTGCTTCC

SDM FWD TGCATTGTTTcCTGTCCTGAAGACGAGC

SDM REV GGGGGTGCCTGCATCTGC

MIN6 cells were seeded at approximately 2.5E05 cells/cm^2 into a 48-well plate. The day after passaging into the 48-well plate, cells were co-transfected with 250ng of experimental firefly luciferase vector pGL4.23 containing the alt or ref allele in the forward direction or an empty pGL4.23 vector, and 15ng pRL-SV40 Renilla luciferase vector (Promega) using the Lipofectamine 3000 reagent. Cells were fed culture media and stimulated where applicable 24 hours post-transfection. Dexamethasone (Sigma) was added to the culture media for dexamethasone stimulation. Cells were lysed 48 hours post transfection and assayed using the Dual-Luciferase Reporter system (Promega). Firefly activity was normalized to Renilla activity and normalized results were expressed as fold change compared to the luciferase activity of the empty vector. A two-sided t-test was used to compare the luciferase activity between the two alleles in each orientation.

## Acknowledgements

This work was supported by DK114650, DK122607, and DK120429 to K.G.

## Author contributions

K.J.G conceived of and supervised the research in this study; K.J.G, A.A and M.O. wrote the manuscript and performed data analyses; J.C., P.B. and E.B. performed data analyses; M.O. performed genomic experiments; M.O., A.P. and S.D. performed reporter experiments.

## Data availability

Processed data and annotations will be made available in https://www.diabetesepigenome.org upon publication, and raw data will be deposited in GEO and dbGAP.

