## Supplementary Figures for "Glucocorticoid signaling in pancreatic islets modulates gene regulatory programs and genetic risk of type 2 diabetes"

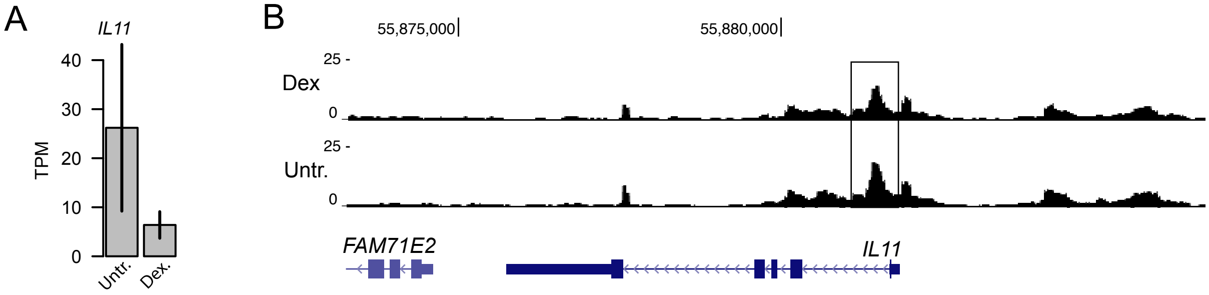


**Supplemental Figure 1. Glucocorticoid treatment downregulates IL11 activity.** (A) Expression level of IL11 in untreated and glucorticoid-treated islets. TPM = transcripts per million. Values represent mean and standard error. (B) Genome browser track showing RPKM normalized ATAC-seq signal in untreated and glucocorticoid treated islets at the IL11 locus. Chromatin site with reduced activity upon glucocorticoid treatment is highlighted.

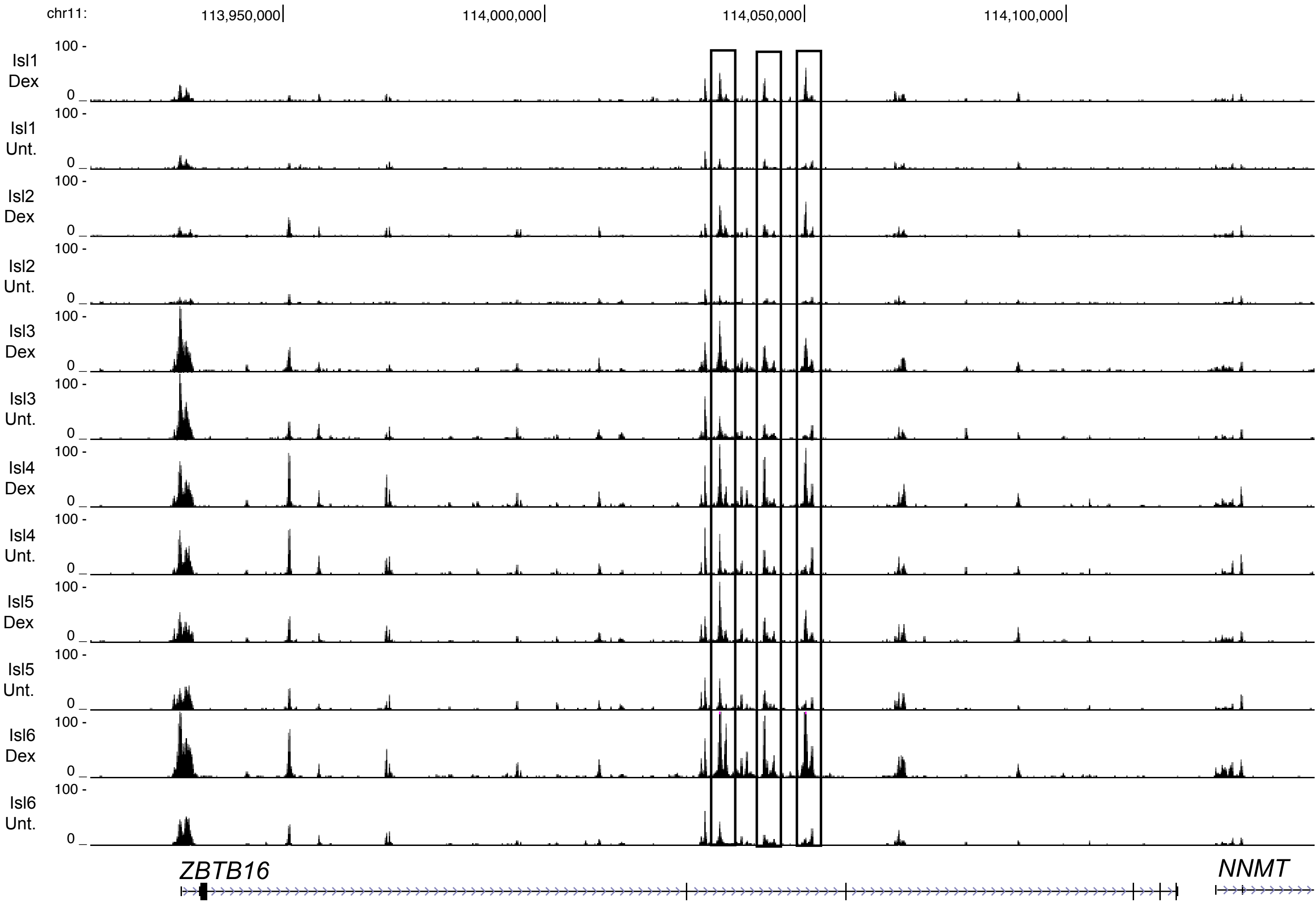


**Supplemental Figure 2. Islet chromatin accessibility at ZBTB16.** RPKM normalized ATAC-seq signal for each individual islet sample in glucocorticoid treated and untreated islets. Sites with differences in chromatin accessibility across conditions are highlighted.

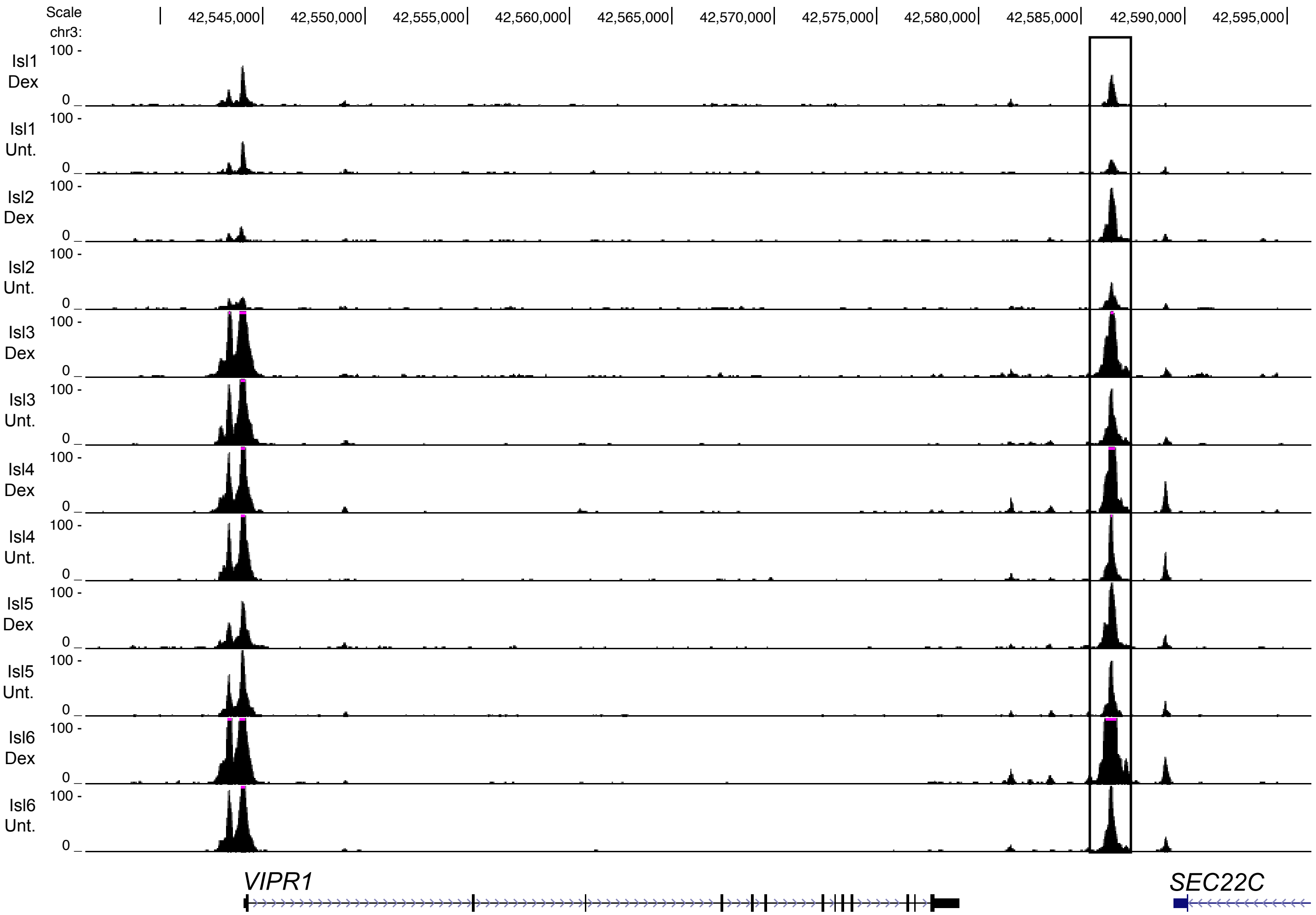


**Supplemental Figure 3. Islet chromatin accessibility at VIPR1.** RPKM normalized ATAC-seq signal for each individual islet sample in glucocorticoid treated and untreated islets. Sites with differences in chromatin accessibility across conditions are highlighted.

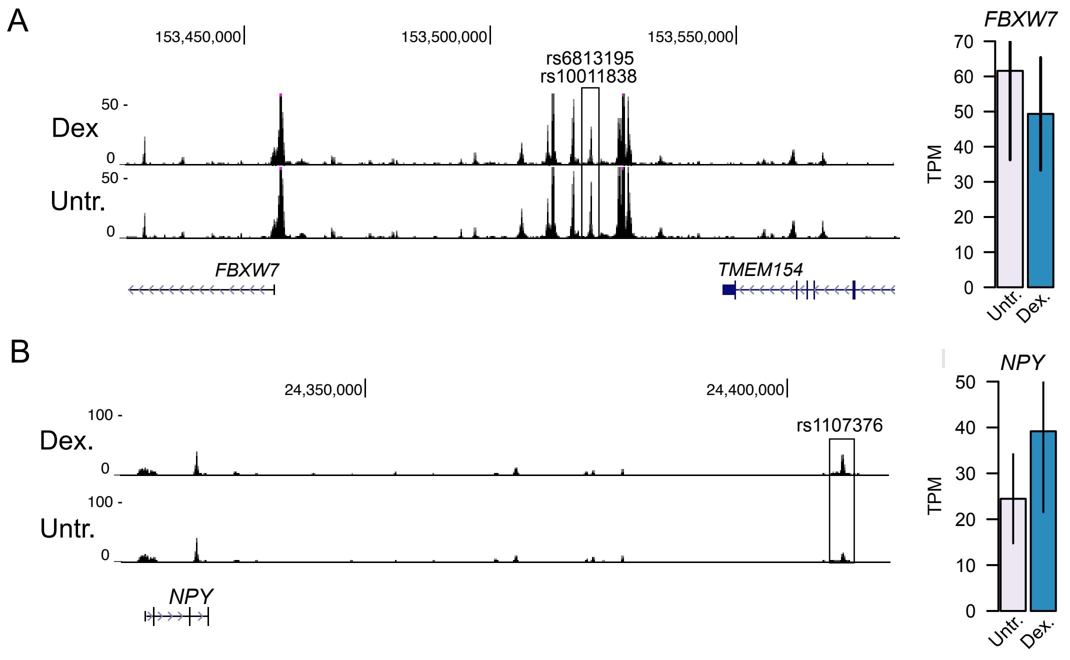


**Supplemental Figure 4. T2D-associated variants in differentially accessible sites.** (A) Multiple variants at the FBXW7/TMEM154 locus mapped in a site with decreased activity and FBXW7 had decreased expression in glucocorticoid stimulation. (B) A variant at the NPY locus mapped in a site with increased activity and NPY had increased expression in glucocorticoid stimulation
